# Divergent host innate immune response to the smooth-to-rough *M. abscessus* adaptation to chronic infection

**DOI:** 10.1101/2023.05.15.540822

**Authors:** Emily A Wheeler, Patricia M. Lenhart-Pendergrass, Noel M. Rysavy, Katie Poch, Silvia Caceres, Kara M. Calhoun, Karina Serban, Jerry A. Nick, Kenneth C. Malcolm

## Abstract

*Mycobacterium abscessus* is a nontuberculous mycobacterium emerging as a significant pathogen for individuals with chronic lung disease, including cystic fibrosis and chronic obstructive pulmonary disease. Current therapeutics have poor efficacy. New strategies of bacterial control based on host defenses are appealing, but anti-mycobacterial immune mechanisms are poorly understood and are complicated by the appearance of smooth and rough morphotypes with distinct host responses. We explored the role of the complement system in the clearance of *M. abscessus* morphotypes by neutrophils, an abundant cell in these infections. *M. abscessus* opsonized with plasma from healthy individuals promoted greater killing by neutrophils compared to opsonization in heat-inactivated plasma. Rough clinical isolates were more resistant to complement but were still efficiently killed. Complement C3 associated strongly with the smooth morphotype while mannose-binding lectin 2 was associated with the rough morphotype. M. abscessus killing was dependent on C3, but not on C1q or Factor B; furthermore, competition of mannose-binding lectin 2 binding with mannan or N-acetyl-glucosamine during opsonization did not inhibit killing. These data suggest that *M. abscessus* does not canonically activate complement through the classical, alternative, or lectin pathways. Complement-mediated killing was dependent on IgG and IgM for smooth and on IgG for rough *M. abscessus*. Both morphotypes were recognized by Complement Receptor 3 (CD11b), but not CR1 (CD35), and in a carbohydrate- and calcium-dependent manner. These data suggest the smooth-to-rough adaptation changes complement recognition of *M. abscessus* and that complement is an important factor for *M. abscessus* infection.

## Introduction

*Mycobacterium abscessus* (*Mab*) is a nontuberculous mycobacteria (NTM) and a significant pathogen in individuals with chronic pulmonary diseases, such as cystic fibrosis, non-CF bronchiectasis and chronic obstructive pulmonary disease (1, 2), as well as soft tissue infections (3). Subclinical infections of the lung may take years to manifest as chronic pulmonary disease (4, 5). *Mab* infection is recalcitrant to most current therapies, which involve extended multi-drug regimens often with poor tolerance and serious side effects (6). Infection is initiated by environmental exposure or person-to-person transmission (7, 8), and is associated with neutrophilic inflammation. Environmental strains have a specific smooth colony morphology when grown on mycobacterial solid media. The smooth morphotype has characteristics that may promote establishing colonization, such as biofilm formation and sliding motility (9). However, the smooth morphotype can have reduced virulence and is less often associated with chronic disease. The smooth morphotype can transition to a rough colony morphotype in vivo. The rough morphotype is associated strongly with increased virulence and immunogenicity, pulmonary disease, and chronic infection (9-15), but mechanisms of rough *Mab* virulence are not well understood. These observations suggest that host factors are insufficient to control infection in at-risk individuals and may promote adaptation to the virulent phenotype. However, little is known of the host factors that control infections in healthy individuals or predispose subjects to *Mab* infection.

Components of the cell envelope of *Mab* are the major determinants that distinguishes the smooth and rough morphotypes. The difference in morphologies is almost entirely due to the loss of glycopeptidolipid (GPL) from the cell wall of the smooth morphotype (9). The lack of GPL on the rough morphotype leads to the surface exposure of mycobacterial-specific molecules such as phosphatidylinositol mannosides (PIMs) and lipoproteins and increased immune cell activation (9, 16, 17). Some details of enhanced virulence with rough *Mab* infection are coming into focus, including enhanced cytokine production and inhibition of autophagy in macrophages infected with rough *Mab* (9, 16-19). However, in neutrophils, cytokine production is similar between the morphotypes, and autophagy does not appear to be involved (20, 21).

Plasma contains pattern-recognition molecules capable of binding and controlling pathogens. These opsonins converge on the activation of the complement pathways of innate immunity. Canonical complement activation consists of the Classical Pathway (CP), the Lectin Pathway (LP), and the Alternative Pathway (AP). The CP is initiated by IgG and IgM binding to pathogens leading to the binding of the C1 complex, a multi-protein protease complex consisting of C1q, C1r, and C1s. The LP is activated by binding of mannose-binding lectin-2 (MBL2) to mannose and other carbohydrates, including N-acetylglucosamine (GlcNAc), and is complexed with MBL2-associated serine proteases (MASPs) (22). The AP uses Factor B in an amplification mechanism to enhance the activity of C3 convertases, which are the common protease activities of all three pathways and cleave C3 to reactive C3b, and further to iC3b. Covalent association of the C3 fragments C3b and iC3b mark pathogens for recognition by the complement receptors CR1, CR3, and CR4 on neutrophils. In addition, specific antibodies can opsonize pathogens for recognition via Fc receptors, although lower affinity non-immune, or natural, antibodies can also act with complement to promote pathogen clearance (23-26).

The role of plasma components to control NTM infections is not well understood. We recently showed that *M. avium* killing by neutrophils is enhanced by plasma complement C3 and IgM (26). To extend these findings to rapidly growing NTM we explored the role of complement and immunoglobulins on killing of *Mab* morphotypes by neutrophils.

## Results

### Plasma components enhance Mab killing by neutrophils

We have previously shown that in the absence of complement *Mab* is poorly recognized or killed by human neutrophils (20, 21, 27). The role of complement as an opsonin was determined by measuring *Mab* survival in the presence of human neutrophils. We compared opsonization with naïve, complement-containing whole plasma (WP) and heat-inactivated plasma (HIP), which is devoid of complement activity. Opsonization of both smooth and rough *Mab* with WP led to rapid and efficient killing by neutrophils (**Fig 1A, B**). Killing with WP opsonization was more robust than with HIP opsonization (**Fig 1C**). A smaller difference between killing in the presence of WP and HIP was observed seen for the rough morphotype of *Mab*, and the difference was significantly different between morphotypes (two-way ANOVA, P=1.5 × 10^−4^). Mock opsonization with BSA resulted in less neutrophil killing than opsonization with WP or HIP (**Fig 1D**). Efficient killing occurred at plasma concentration as low as 5% (**Fig 1E, F**). Reaction of WP or HIP in the absence of neutrophils did not affect *Mab* survival over the times used in these studies (**Suppl Fig 1**). The heat-labile nature of plasma components suggests a role for complement opsonization in the recognition and clearance of *Mab*.

**Fig 1.**
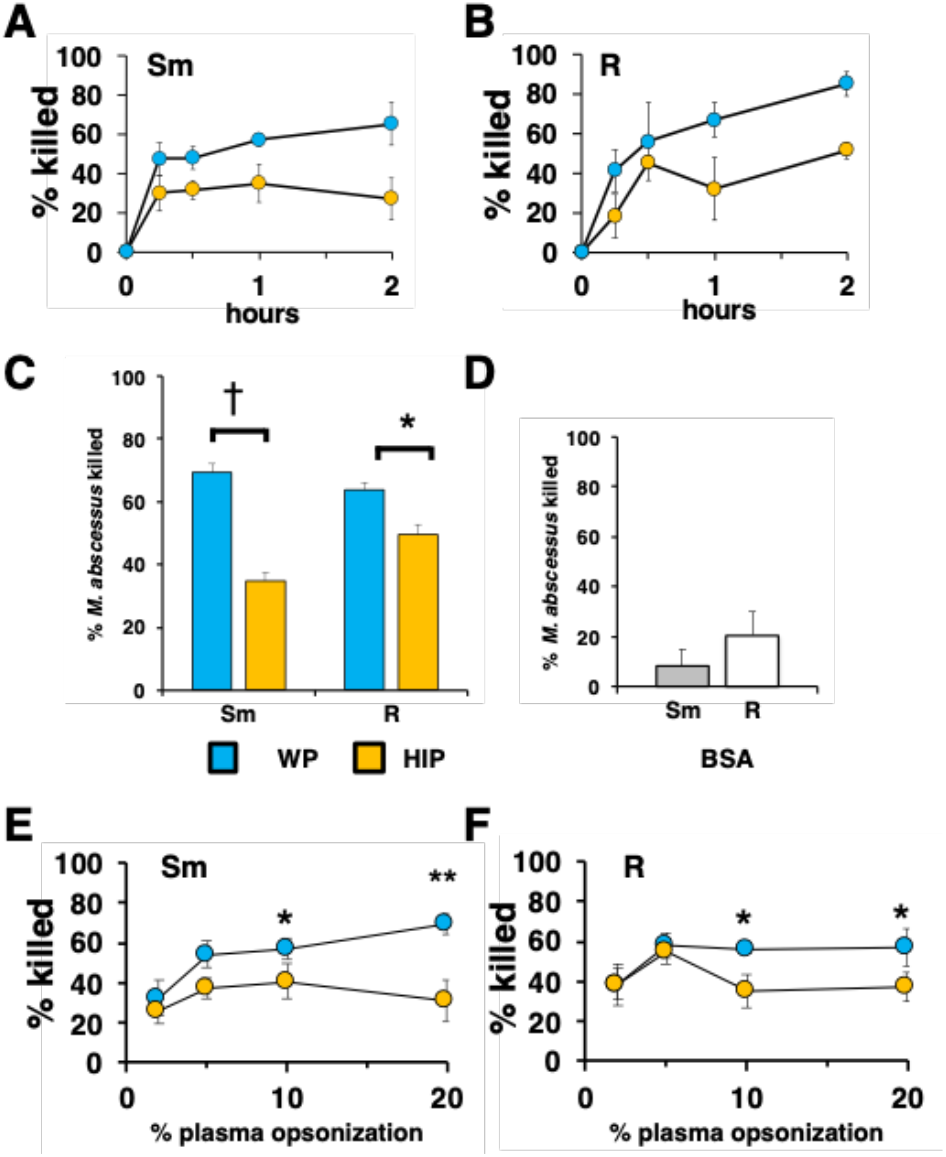
Whole plasma enhances killing of *Mab*. **A**, smooth (Sm) *Mab* and **B**, rough (R) *Mab* opsonized with whole plasma (WP; blue) or heat-inactivated plasma (HIP; orange) were added to human neutrophils at an MOI of 1 for the indicated times and killing determined; n=3. **C**, smooth and rough *Mab* were opsonized in WP or HIP; n=37; †, P = 1.2 × 10^−11^, 8, 1.8×10^−5^), or **D**, incubated with BSA (n=4), and killing determined after 1 hr incubation with neutrophils. **E**, smooth, and **F**, rough *Mab* opsonized with the indicated percentage of WP or HIP and neutrophil killing determined after 1 hr incubation; n=5-6; *, P<0.05; **, P<0.01.

The need for pre-opsonization of *Mab*, as performed in the above experiments, was also addressed to determine if the effect of complement opsonization was influenced by the presence of neutrophils. When plasma was introduced to the neutrophils just before non-opsonized *Mab* was added, killing of smooth *Mab* was reduced, but killing of rough *Mab* was not changed, when compared to pre-opsonized *Mab* (**Suppl Fig 2**). Reduced smooth *Mab* killing with simultaneous WP in the presence of neutrophils may reflect altered kinetics of opsonization compared to the rough morphotype or the activity of neutrophil-derived negative regulatory factors. Drawing blood into different types of collection tubes can affect plasma components (28). Therefore, we compared the killing ability of serum and of plasmas from blood drawn into citrate, heparin, and EDTA. Citrate and EDTA WP were re-calcified and clotted. Killing was similar in all conditions (**Suppl Fig 3**), suggesting that plasma source is not a major constraint in measuring complement function.

### Smooth and rough morphotypes deposit distinct complement factors

The plasma protein components associated with *Mab* in the various opsonization conditions were explored by immunoblotting. *Mab* was opsonized with WP, washed, and *Mab*-associated proteins extracted in SDS sample buffer. Blots were probed with antibodies to iC3b, C1q, MBL2, IgG and IgA (**Fig 2A-C**). C3 fragments associated with smooth *Mab*, but little MBL2 and C1q deposition was observed. C3-positive bands represent the C3 fragments, including those covalently bound to *Mab* moieties through the reactive ester, with major bands at 70, 120, and a group at 260 kDa. Rough *Mab* association with C3 was distinct from that of smooth *Mab*; little of the major 70 and 130 kDa bands were observed, but the 260 kDa bands were at similar levels as for smooth *Mab*. C1q deposition was weak. MBL2 association was more pronounced in rough *Mab*. The immunoglobulin classes IgG, IgA, and IgM bound to the *Mab* morphotypes (**Fig 2B, C)**. These data indicate morphotype-specific activation of C3 and MBL2.

**Fig 2.**
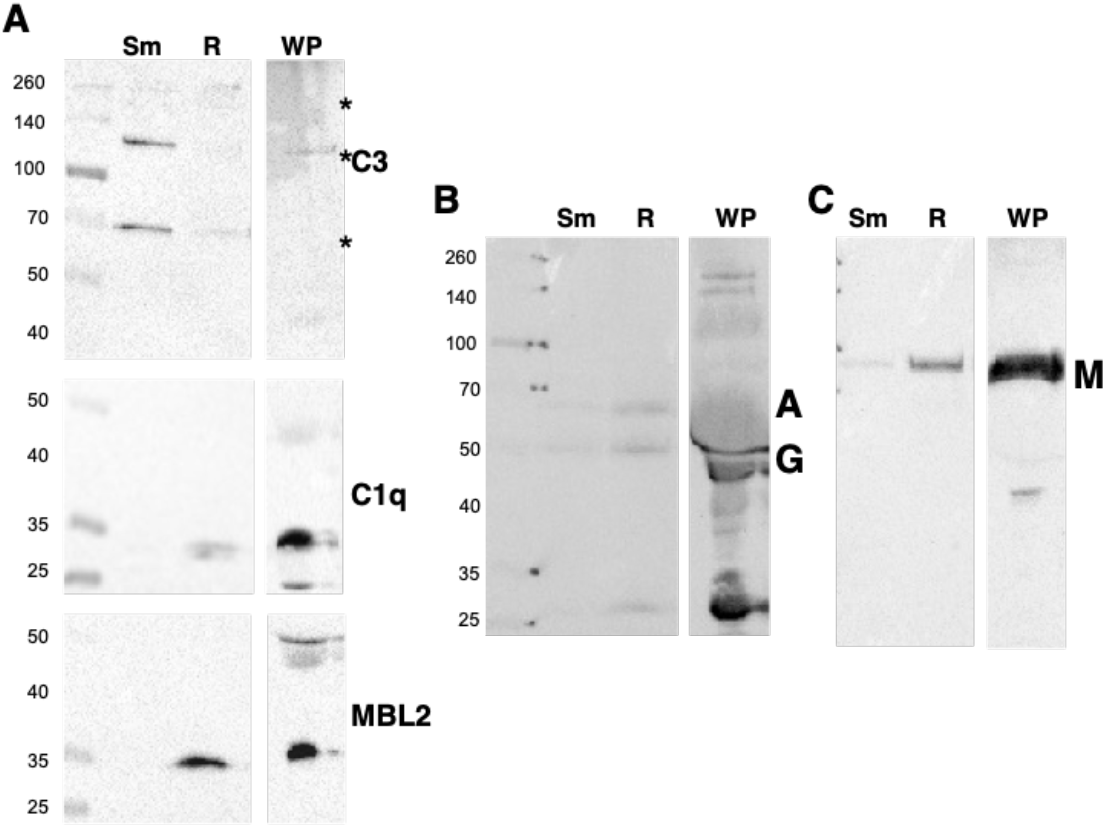
Whole plasma differently opsonizes *Mab* morphotypes. **A**, smooth (Sm) and rough (R) *Mab* opsonized with whole plasma were washed, proteins separated by SDS-PAGE, and deposited C3, C1q, and MBL2 were detected. A sample of WP was run as a positive control; * indicates C3 fragments. **B**, detection of IgG (G) and IgA (A). **C**, detection of IgM (M). The results are representative of over 10 separate experiments. Molecular weight markers are shown to the left of each blot.

### Smooth and rough clinical isolate opsonization and killing

To determine if morphotype differences between C3 and MBL deposition was a general phenomenon, we measured complement deposition on genetically distinct CF clinical smooth and rough isolates from people with CF (29). Like the Type strains, C3 deposition of the 70 and 120 kDa bands was predominant on the smooth strains. MBL2 deposition was enhanced on the rough strains in the isogenic pairs (ATCC and CF016) (**Fig 3A**). C1q was minimally detected. However, MBL2 deposition was variable among isolates where we observed more deposition of MBL2 on CF0123, a smooth strain, then on the genetically related rough strain CF0017-R. The effect on killing after opsonization of these clinical isolates was determined. Smooth isolates were more susceptible to heat-sensitive plasma components than rough isolates (**Fig 3B**), either individually or in composite (**Suppl Fig 4**). These data suggest that rough isolates are less sensitive to complement than smooth isolates, although exceptions exist.

**Fig 3.**
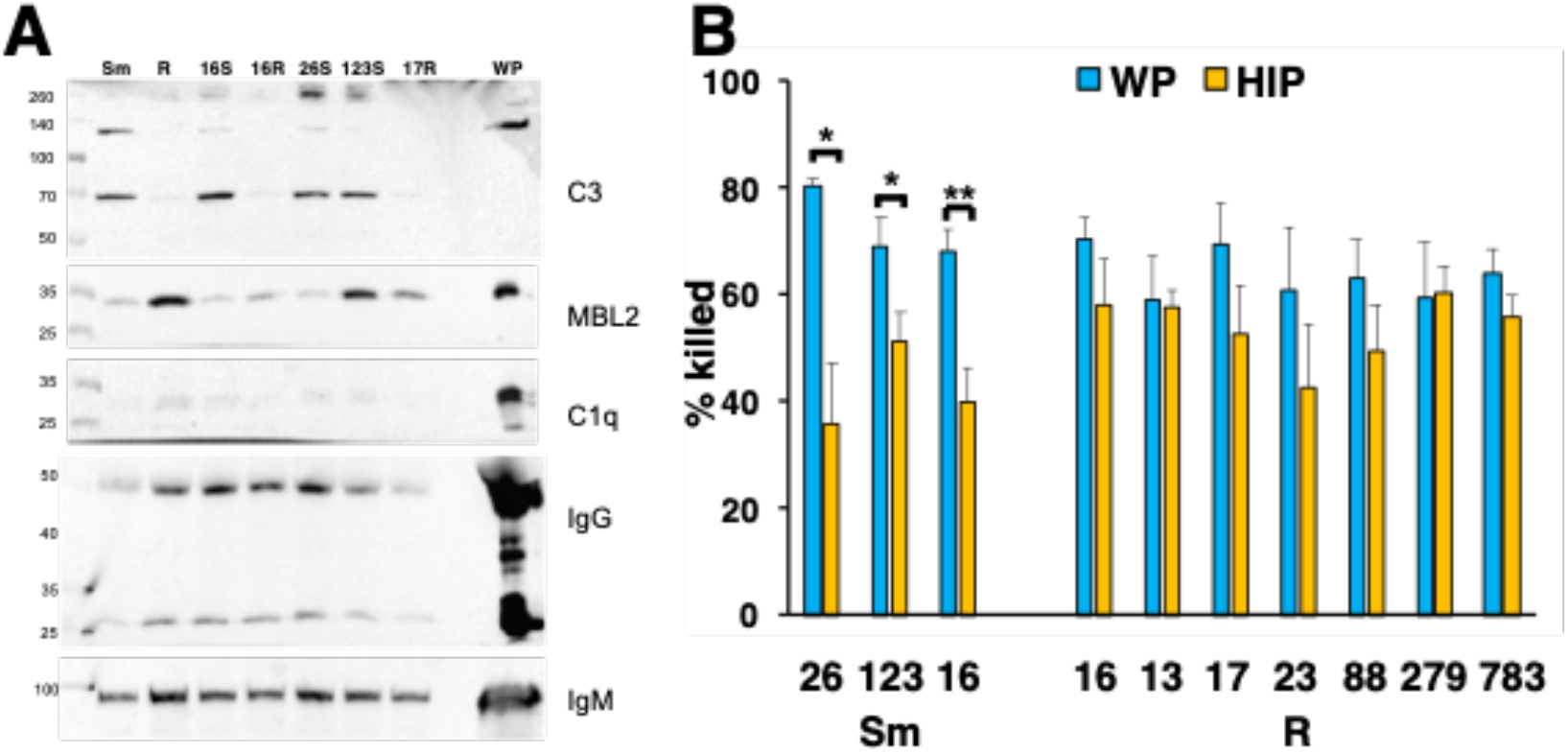
Killing of smooth *Mab* clinical isolates is more dependent on complement than rough *Mab* isolates. **A**, smooth (S) and rough (R) *Mab* clinical isolates opsonized with WP were washed, proteins separated by SDS-PAGE, and deposited C3, C1q, MBL2, IgG and IgM were detected. A 1:20 dilution of WP was run as a positive control. S and R designate the morphotype. *, P<0.05; **P<0.01. **B**, smooth and rough *Mab* clinical isolates were opsonized with WP (blue) or HIP (orange) were added to human neutrophils at an MOI of 1 for 1 hr and killing determined; n = 4-11 per isolate.

### C3 promotes neutrophil Mab killing

The activation of C3 is common to the three major pathways of complement activation. Opsonization of *Mab* with serum depleted of C3 led to a reduction in killing of both morphotypes (**Fig 4A, B**), but the inhibition was greater for smooth *Mab*. The observations that rough *Mab* killing was dependent on C3, but that little C3 deposition was observed, could be from reduced C3 activation or from loss of reacted C3 from the cell surface (30, 31). To address this, we measured residual C3 in the supernatant after the opsonization reaction. Incubation of WP with rough *Mab* resulted in the preservation of C3 in the post-reaction supernatant of similar size (∼130 kDa) as unreacted C3 in the WP (**Fig 4C**), a finding consistent with reduced C3 activation. The post-opsonization supernatant of smooth *Mab* had minimal unreacted C3 indicating efficient reaction and deposition on its surface. These data confirm a role for C3 as a smooth *Mab* opsonin, but with reduced C3 activation by rough *Mab*.

**Fig 4.**
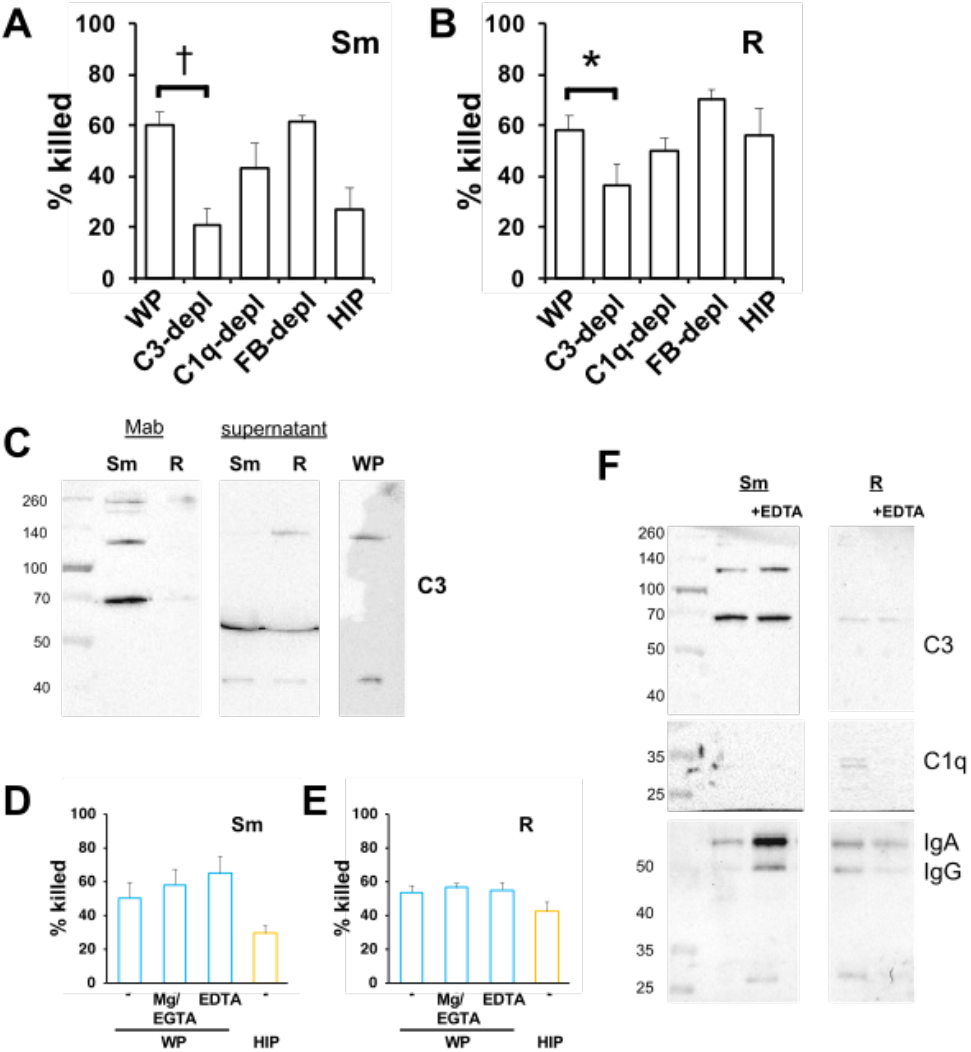
Opsonized killing of *Mab* requires C3, but is independent of CP and AP. **A**, smooth *Mab* and **B**, rough *Mab* opsonized with serum, sera depleted of C3, C1q, and Factor B, or HIP were added to human neutrophils at an MOI of 1 for 1 hr and killing determined; n = 6-8. *, P<0.05; **P<0.01. **C**, smooth and rough *Mab* were opsonized with WP and the supernatants were retained. The washed cellular pellet and supernatant proteins were separated by SDS-PAGE, and deposited C3 was detected. A 1:20 dilution of WP was run as a positive control; representative of 3 independent experiments. **D**, smooth, and **E**, rough *Mab* were opsonized with WP or HIP alone, or in the presence of Mg2+/EDTA or EGTA before adding to neutrophils. Killing was determined after 1 hr incubation; n = 6. **F**, smooth and rough *Mab* were opsonized in WP or WP + EDTA, washed, and deposited C3, C1q, IgG, and IgA were detected; n = 3. *, P<0.05; †, P<0.001.

### Opsonization occurs in the absence of the CP and AP

To determine the role of the CP and AP on C3 activation we opsonized *Mab* with serum depleted of C1q and Factor B, mediators of the CP and AP, respectively. Depletion of C1q or Factor B did not reduce killing, indicating that opsonization occurred independently of the CP and AP (**Fig 4A, B**). To confirm these findings, *Mab* was opsonized with WP in the presence of Mg^2+^/EGTA, a condition that allows only AP activation, or EDTA to chelate Ca^2+^ and Mg^2+^ and inhibit activation of all three complement pathways (**Fig 4D, E**). Interestingly, these conditions did not reduce *Mab* killing, notably in light of the association of MBL2 with rough *M. abscessus*. Consistent with the killing data opsonization in the presence of EDTA did not affect deposition of C3 on *Mab* (**Fig 4F**) or the deposition of IgG or IgA. These data suggest that canonical activation of the CP, LP, and AP are not essential for WP opsonization.

### Competition of MBL2 with mannan and GlcNAc does not alter killing of Mab

To address the role of MBL2 in *Mab* opsonization, we performed the opsonization reaction in the presence of competitive soluble mannan (an α-linked mannose polymer) and GlcNAc (N-acetyl-D-glucosamine), two agents that bind MBL2 to activate the MASP2-dependent C3-convertase. Opsonization in the presence of mannan or GlcNAc did not affect killing of *Mab* (**Fig 5A**). In the same conditions, association of C3 and MBL2 was measured. Although mannan led to increased MBL2 association with both morphotypes C3 deposition was not affected (**Fig 5B**). In contrast, GlcNAc reduced association with MBL2 with little effect on C3 deposition. Association with IgG and IgM did not change in either condition. These data suggest that C3 fixation can occur independently of MBL2.

**Fig 5.**
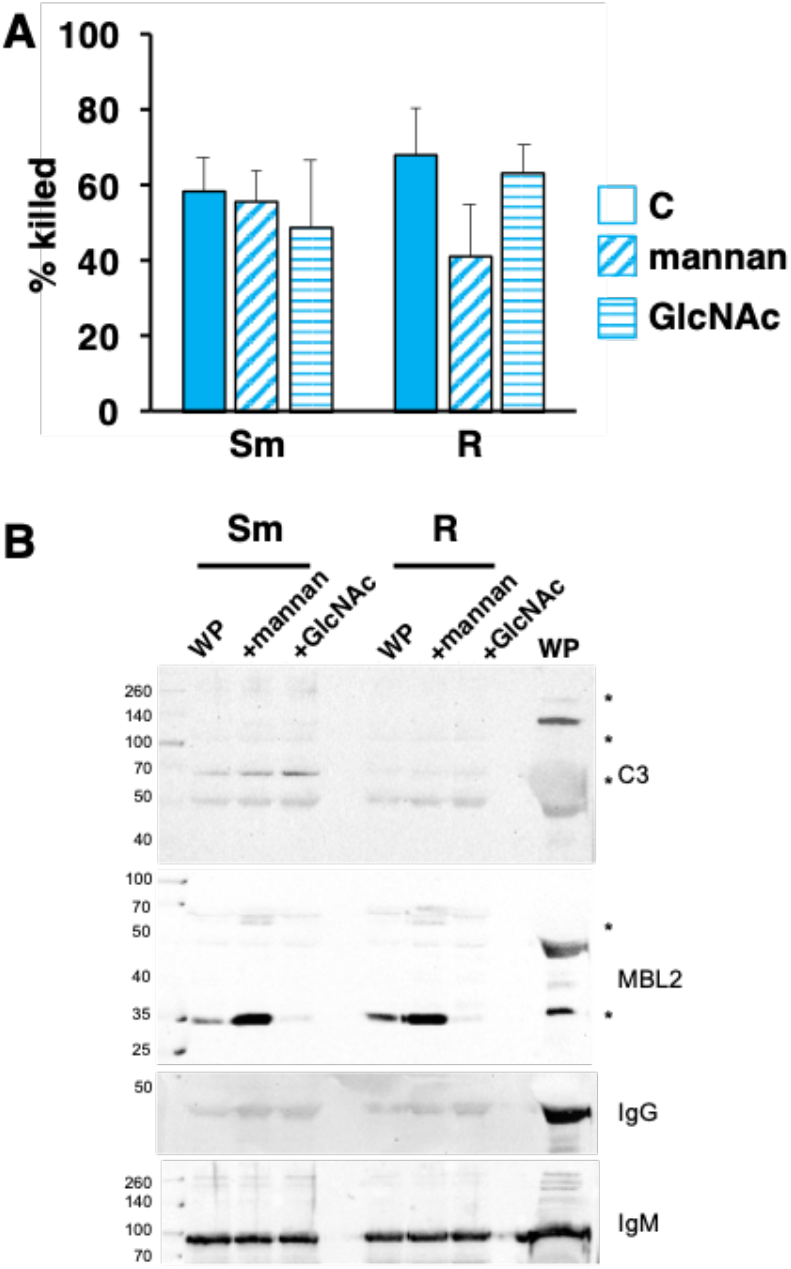
Opsonized killing of *Mab* is not inhibited by MBL2 ligands. **A**, smooth and rough *Mab* opsonized with WP alone (solid) or in the presence of mannan (diagonal stripes; 1 mg/ml) or GlcNAc (horizontal stripes; 100 mM) were added to human neutrophils at an MOI of 1 for 1 hr and ki l l ing determined; n= 5. **B**, *Mab* opsonized as in A were washed, proteins separated by SDS-PAGE, and deposited C3, MBL2, IgG, and IgM were detected. A 1:20 dilution of WP was run as a positive control.

### Smooth and rough Mab require distinct Ig recognition to mediate opsonization

In addition to activation of the CP, antibodies can act as opsonins to mediate phagocytosis and killing in the absence of complement. The role of immunoglobulins in the recognition and killing of *Mab* was explored using depleted plasmas. We previously observed a role of IgM in complement-dependent neutrophil *M. avium* killing (26). To explore the role of IgM in opsonization of smooth *Mab*, we depleted IgM from plasma with anti-IgM. WP depleted of IgM reduced killing of smooth, but not rough, *Mab* (**Fig 6A, B**). Depletion of IgM with anti-IgM was previously confirmed (26). Consistent with IgM lacking a direct opsonizing effect, IgM depletion did not affect killing when *Mab* were opsonized with HIP. These data indicate that IgM is a complement-dependent opsonin of only smooth *Mab*.

**Fig 6.**
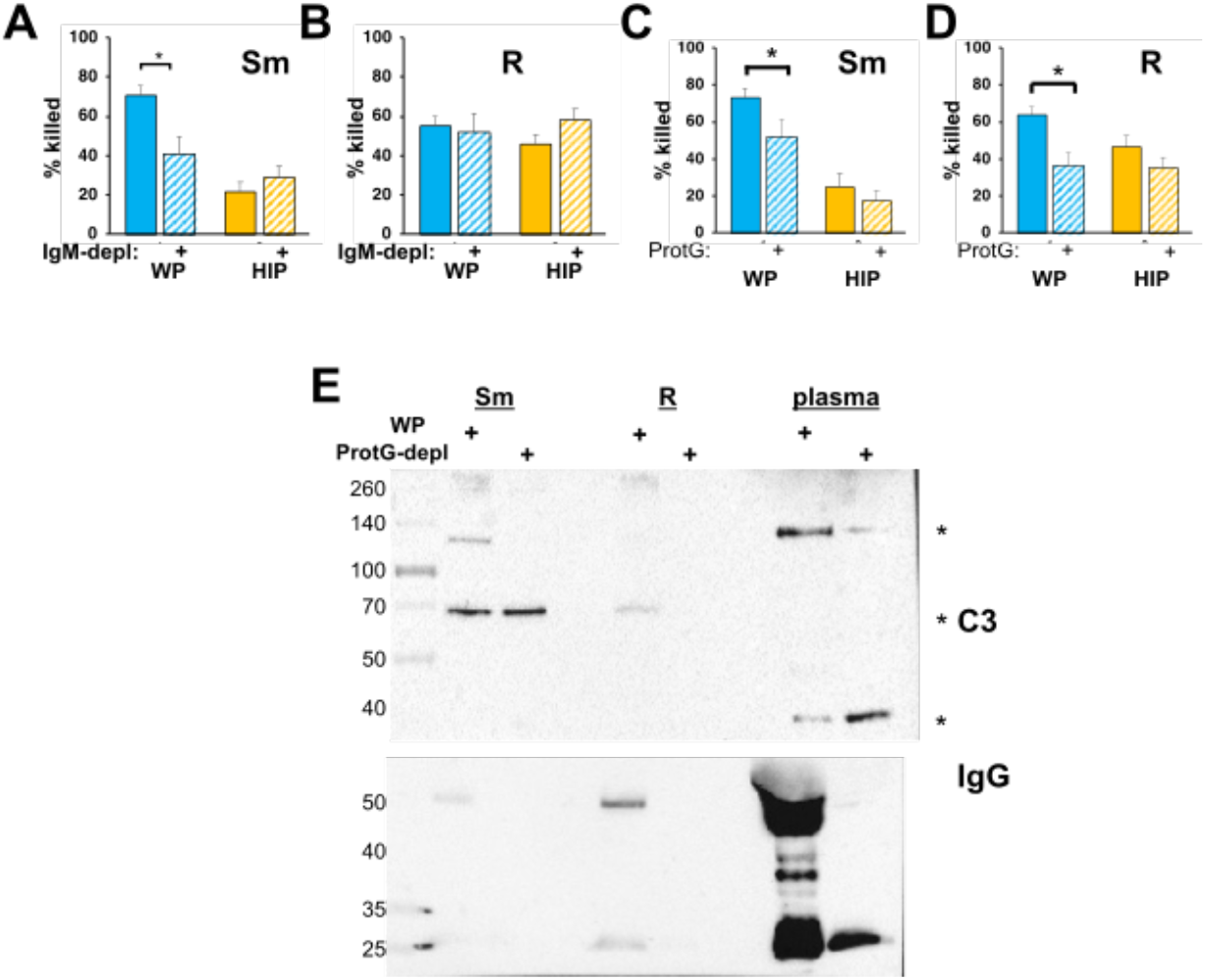
Immunoglobulins are required for killing of *Mab* opsonized in WP. **A**, smooth and **B**, rough *Mab* opsonized with WP or HIP alone (solid) or in WP or HIP depleted of IgM with anti-IgM agarose (striped bars) were added to human neutrophils at an MOI of 1 for 1 hr and killing determined. **C**, smooth and **D**, rough *Mab* opsonized with WP or HIP alone (solid) or in WP or HIP depleted of IgG with Protein G Sepharose (striped bars) were added to human neutrophils at an MOI of 1 for 1 hr and killing determined; n= 5-10 independent experiments. **E**, *Mab* opsonized as in Fig 2 were washed, proteins separated by SDS-PAGE, and deposited C3 and IgG were detected. A 1:20 dilution of WP or IgG-depleted WP (plasma) were run as positive controls; * indicates C3 fragments.; n= 5. *, P<0.05.

To understand the role of IgG, the four subclasses of IgG were selectively depleted using Protein G. Opsonization in Protein G-depleted WP reduced *Mab* killing (**Fig 6C, D**). The incomplete reduction of killing of smooth *Mab* in Protein G-depleted WP suggests additional factors, such as IgM, are involved. Depletion of IgG from HIP had no effect on *Mab* killing, suggesting a lack of direct effect of IgG opsonization on killing. Consistent with the killing data, C3 deposition was reduced in IgG-depleted serum (**Fig 6E**), as was association of IgG with *Mab*. These data suggest that the opsonins of *Mab* include IgG and IgM for smooth Mab and IgG for rough Mab, and that IgG and IgM are necessary but not sufficient for optimal *Mab* killing in the presence of WP. Similar results were obtained using depletion with Protein A, which binds IgG (with the exception of the IgG3 subclass) and IgM, and to a lesser extent IgA (**Suppl Fig 5**).

### Receptor recognition of opsonized Mab

Receptors for C3 include the integrin family of adhesion receptors, which require divalent cations for efficient binding (32). Reduction of Ca^2+^ using EGTA and of Ca^2+^ and Mg^2+^ with EDTA during the killing assay resulted in decreased killing of *Mab* opsonized with WP and HIP (**Fig 7A, B**), suggesting the importance of calcium-dependent recognition of complement and non-complement opsonized *Mab* by neutrophils. To specifically address the role of neutrophil complement receptors, we used blocking antibodies to CR1 (CD35), which binds C3b, and to CR3 (CD11b/CD18), which binds iC3b. Anti-CD11b was partially effective at blocking killing (**Fig 7C, D**), anti-CD35 had no effect on killing, and the combination of anti-CD35 and -CD11 (24) was similar to anti-CD11b alone. Inhibition by anti-CD11b did not reach the reduced levels seen for HIP-opsonized *Mab*. These data indicate that CR3 has a role in the recognition of *Mab* opsonized with WP, but that cations are important for complement-dependent and -independent recognition by neutrophils.

**Fig 7.**
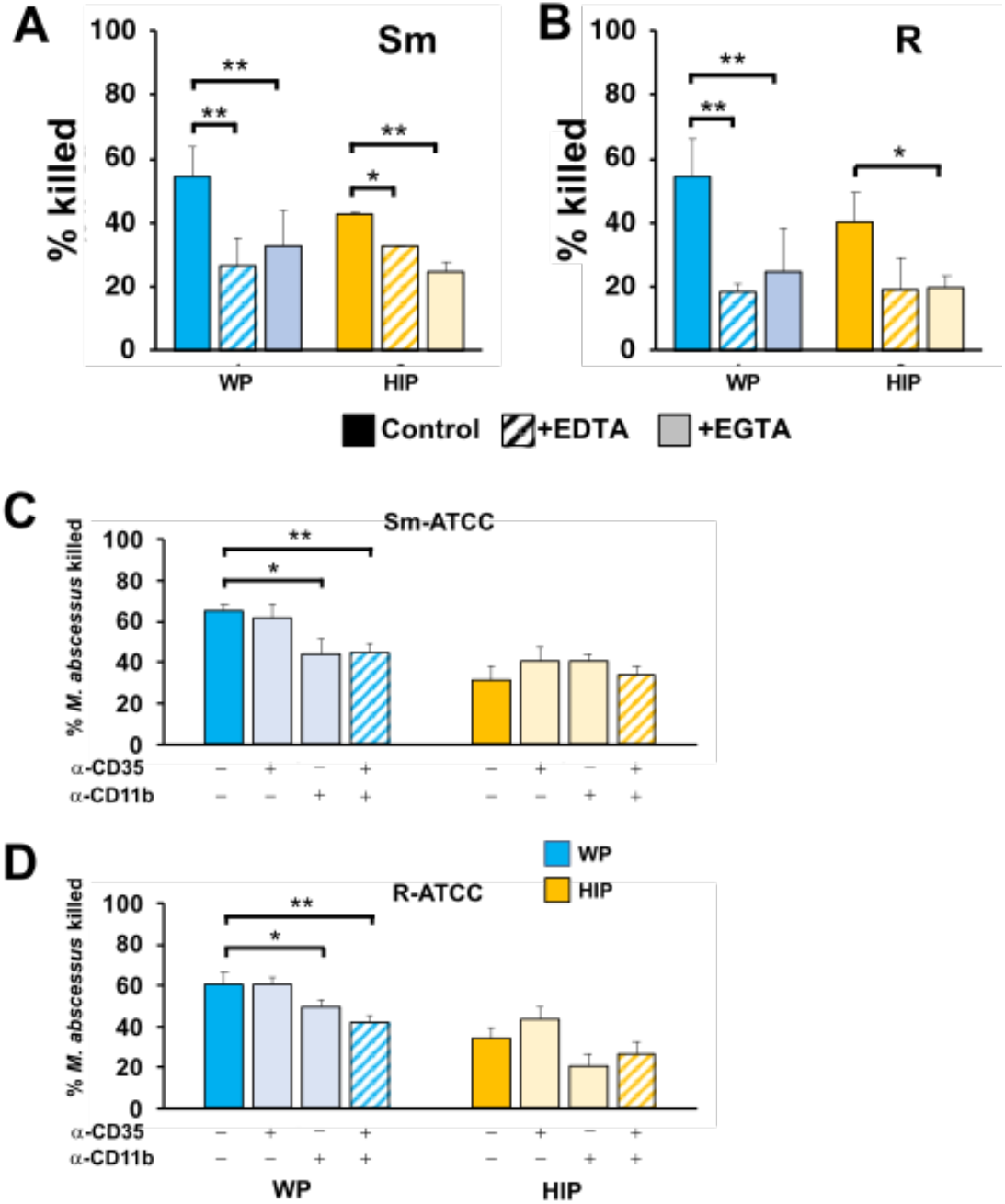
CR3/CD11b and cations play independent roles in killing of opsonized *Mab*. **A**, smooth and **B**, rough *Mab* opsonized with WP or HIP (solid) were added to human neutrophils in the presence of EDTA (striped bars) or EGTA (light shading) at an MOI of 1 for 1 hr and killing determined; n= 4-6. **C**, smooth and **D**, rough *Mab* opsonized with WP or HIP (solid) were added to human neutrophils pre-incubated with anti-CD35 (CR1) or anti-CD11b (CR3), as indicated, and killing determined; n= 5-6. *, P<0.05; **P<0.01.

### Carbohydrates block Mab killing

To further define the recognition receptors on neutrophils we explored the ability of free carbohydrates that mimic surface moieties to affect *Mab* killing. GlcNAc, GalNAc (N-acetyl-D-galactosamine), and mannose are sugars found on mycobacterial cell walls (33). These sugars also activate complement (22, 34) and interact with host and pathogen receptors (35-37). Incubation of neutrophils with either GlcNAc or GalNAc reduced *Mab* killing after the bacteria was opsonized with either WP or HIP (**Fig 8A, B**).

**Fig 8.**
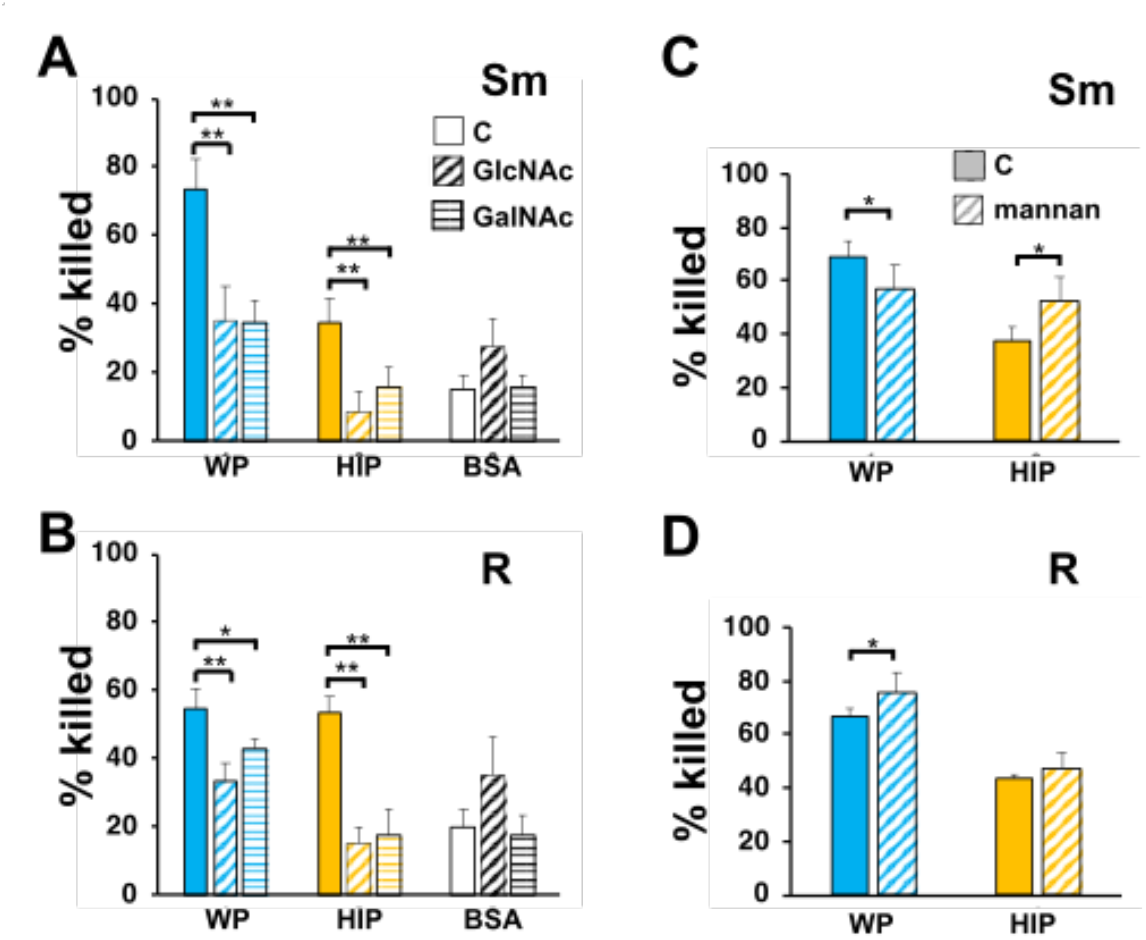
Inhibition of *Mab* killing by N-acetylated sugars. **A**, smooth and **B**, rough *Mab* opsonized with WP or HIP alone (solid) or incubated with BSA were added to human neutrophils in the presence of GlcNAc (diagonal stripes; 100 mM) or GalNAc (horizontal stripes; 100 mM) for 1 hr and killing determined; n=5. **C**, smooth and **D**, rough *Mab* opsonized with WP or HIP alone (solid) were added to human neutrophils in the presence of mannan (diagonal stripes; 1 mg/ml) for 1 hr and killing determined; n=6 *, P<0.05; **P<0.01.

They had no effect on the minimal killing of *Mab* mock opsonized with BSA. Incubation with mannan did not consistently affect *Mab* killing (**Fig 8C, D**). Minimal inhibition of smooth *Mab* opsonized with WP was seen, but a small, enhanced killing of HIP-opsonized smooth *Mab* and WP-opsonized rough *Mab* were also observed. Therefore, recognition of N-acetylated sugars, but not mannose, is important for neutrophil killing of *Mab*, although these carbohydrate interactions likely involve both complement-dependent and independent mechanisms.

## Discussion

Host mechanisms to contain NTM infections are poorly understood. Innate immune cells are implicated in the control of infections, but detailed mechanisms of clearance have not been addressed. Furthermore, most studies have focused on the interaction of myeloid cells with NTM in the absence of other host-derived factors. The uptake and killing by neutrophils of the slow growing NTM *M. avium* was increased in a complement and IgM dependent manner (26, 38). However, the role of complement in the control of rapidly growing NTM has not been explored. Here, we have demonstrated for the first time antibody-dependent complement opsonization of rapidly growing NTM.

Our findings are notably in that the adapted, rough form has acquired mechanisms to evade two different innate immune mechanisms of pathogen control, namely C3 and IgM opsonization, resulting in MBL2 and IgG specificity. This finding suggests that C3 and IgM are in-host selective pressures for adaptation in the diseased lung. However, this change in fitness comes at the expense of promoting MBL2 opsonization and without compromising the effects of IgG. The observations that complement promotes *Mab* clearance and that more severe *Mab* pulmonary disease is associated with a slow rate of conversion to the rough morphotype suggest that a subtle balance exists between adaptation and immune control of the two morphotypes. Support for the growth of specific morphotypes may rely on micro environments that contain different growth conditions or complement and Ig class repertoires. For instance, vascular leakage may enrich for complement in an injured lung compared to a healthy lung (39). Killing of both morphotypes was reduced in C3-depleted serum, suggesting that the minimal C3 deposited, as evident from the 260 kDa bands on rough *Mab*, is still sufficient to drive at least some killing.

Differences in complement activation and function between the morphotypes is further highlighted by clinical isolates. Killing of smooth *Mab* isolates was more dependent on heat-sensitive plasma components than rough *Mab* isolates. These data support the idea that the smooth morphotype may gain a foothold for sustained infection at infection sites that are limited or deficient in complement factors (20, 21), and transition to the more virulent rough morphotype occurs under prolonged selective pressure. A recent report of reduced complement levels in the CF lung support this idea (40)

The dependence of complement activation on antibodies was unexpected. The plasma donors were healthy, and we previously showed minimal anti-*Mab* specific antibodies in healthy individuals (41). Our findings implicate natural antibodies in complement activation, as previously reported (23-26, 42). The antibody specificities required for full opsonization of the morphotypes may reflect changes in cell surface components. Loss of glycopeptidolipid (GPL) that is the hallmark of rough *Mab* is accompanied by exposure of phosphatidylinositol mannosides (PIMs) (16) and overexpression of lipoproteins (17). These exposed molecules may represent novel epitopes for non-immune IgG. Other cell surface composition differences may also be involved (43-45). Differences in host Fc receptor utilization or activation may also confer specificity. Interaction of CR3 (CD11b/CD18) and FcγRIII suggests one such mechanism (46, 47), but the exact mechanism of antibody-complement interaction remains to be seen.

Surprisingly, IgG- and IgM-dependent C3 activation by smooth *Mab* appears to be independent of the CP based on studies conducted in the absence of C1q, minimal C1q deposition on *Mab*, and lack of effect of EDTA to block opsonization. Similarly, the AP does not seem to play a major role as killing and C3 deposition are not inhibited by EDTA and killing is not inhibited in Factor B-depleted serum. A similar independence from CP and AP was found for killing of *M. avium* (26). Reduced MBL2 association with smooth *Mab* and no effect of EDTA on C3 deposition or killing do not support a role for the LP. Furthermore, mannan enhanced MBL2 deposition and GlcNAc reduced MBL2 deposition without altering killing. These data suggest that canonical complement activation are not major drivers of smooth *Mab* killing.

Similar observations as for smooth *Mab* indicate that opsonization of rough *Mab* does not involve CP and AP. The data is less clear for a role of the LP for opsonization of rough *Mab*. Killing or opsonization was not blocked by EDTA, Mg^2+^/EGTA, mannan, or GlcNAc; these agents, with the exception of GlcNAc, also did not reduce MBL2 deposition. As seen with smooth *Mab*, GlcNAc reduced MBL2 deposition without affecting killing. However, rough *Mab* preferentially associates with MBL2. These data suggest that *Mab* killing includes pathways distinct from canonical complement activation. This discrepancy may be explained by direct MBL2 opsonization (48-50), which was shown for M. avium. The surface exposure of PIMs or mannose-capped lipoarabinomannan in rough *Mab* may be the basis of the association of MBL2 with rough *Mab*. In addition, MBL2 can activate C3 in the absence of C2/C4 activation, in a process termed “C2/C4 bypass” (51). The interaction of C3 and MBL2 on rough *Mab* remains unexplained.

Non-canonical plasma opsonins and complement activators could explain the morphotype-specific profiles of C3 activation. Alternative factors may include members of the pentraxin family (Pentraxin 3, C-reactive Protein, and Serum amyloid P) or ficolins (FCNs), all of which are pattern-recognition receptors associated with complement activation (52). Preliminary experiments showed little to no deposition of the pattern recognition receptors FCN1, FCN2, FCN3, or Pentraxin 3 to *Mab* (not shown), but these experiments may have been limited by poor antibody recognition. In addition, binding to ficolins is Ca^2+^-dependent (53), and EDTA did not reduce C3 deposition or killing, suggesting that they are not involved. Further study is warranted to clarify alternative pathways for C3 deposition.

Evidence supports recognition of WP-opsonized *Mab* by CR3, an integrin receptor that plays a major role in the binding and phagocytosis of iC3b-opsonized pathogens (54). Antibody blockade of CR3 reduced *Mab* killing, and inhibition of killing by EGTA and EDTA support a role of integrins. CR3 binds GlcNAc, which may interfere with the interaction of CD11b and FcγRIII (47). However, the profound inhibition of killing by GlcNAc and GalNAc, in both WP- and HIP-opsonized *Mab*, indicate that non-complement lectins are also involved. The greater inhibition by acetylated sugars may also reflect the ability of bacterial lectins to recognize host GlcNAc (36, 37). Non-receptor mycobacterial recognition has also been described (55). These data suggest that both CR3 and non-complement-dependent receptors are important recognition events.

Limitations to this study exist. We were unable to completely define the complement activation pathways. Deposition of C3 on *Mab* in the absence of CP, AP, or LP is unusual. Complement activation is controlled by positive and negative regulators that were not explored. Our studies focused on the role of non-immune plasma. We have previously demonstrated that anti-*Mab* IgG exist in NTM-infected people with CF (41), but the role of specific immunity on opsonization and killing will require further study. Finally, these studies focus on neutrophils; the role of complement for other immune cells working and in vivo effects require further study.

In summary, we have identified C3-dependent killing of *Mab* by neutrophils that require distinct opsonization patterns for the smooth and rough morphotypes. The data indicate that opsonization of *Mab* involves non-canonical complement activation that is dependent on specific natural antibody classes for smooth and rough *Mab* morphotypes. Together, these studies support distinct pathways for morphotype killing by neutrophils, suggest complement deficiency may be a factor in disease risk, and have implications in the role of complement in *Mab* adaptation to the diseased lung.

## Materials and Methods

### Bacterial strains, media, and culture conditions

*M. abscessus* ssp. *abscessus* (*Mab*; ATCC strain 19977) smooth and rough morphotypes were propagated from frozen aliquots in 7H9 broth supplemented with 0.5 g/l bovine albumin fraction V,

0.2 g/l dextrose,

0.3 mg/l catalase (ADC; BD Biosciences), 2% glycerol and 0.05% Tween-80 at 37°C with shaking at 200 rpm for 3-4 days. Clinical isolates were obtained from the Culture, Biorepository, and Coordinating Core of the Colorado Cystic Fibrosis Foundation Research Development Program at National Jewish Health. Washed cultures were sonicated using six one-second bursts of a Fisher Sonic Dismembrator 100 at approximately 4 W output to obtain single cells and smaller aggregates and adjusted to OD_600_ of 1.0 in PBS containing Ca^2+^ and Mg^2+^, corresponding to approximately 3 ×10^8^ cfu/ml (27).

### Study population

Blood was drawn from healthy donors for isolation of neutrophils and plasma. These studies were approved by the Biomedical Research Alliance of New York Institutional Review Board, and written informed consent was obtained from all donors. The study was conducted in accordance with the Declaration of Helsinki.

### Neutrophil isolation

Neutrophils were isolated from healthy volunteers by the plasma Percoll method as previously described (56) from blood drawn into 13 mM sodium citrate. Isolated neutrophils were washed and resuspended in Krebs-Ringer phosphate-buffered dextrose (154 mM NaCl, 5.6 mM KCl, 1.1 mM MgSO_4_, 2.2 mM CaCl_2_, 0.85 mM NaH_2_PO_4_, 2.15 mM Na_2_HPO_4_ and 0.2% dextrose). Cells were confirmed to be >98% pure by visual inspection of cytospins.

### Plasma isolation

Citrated platelet-rich plasma from the neutrophil isolation was centrifuged at 2000 x g for 15 min to obtain platelet-poor plasma and re-calcified with the addition of 40 mM CaCl_2_ to allow clotting. Plasma from EDTA tubes was recalcified with 5 mM CaCl_2_ and clotted as for citrated plasma. Serum was clotted for > 1 hr at room temperature and all plasmas were isolated after centrifuged at 2000 x g for 15 min. Because serum tubes may have components that inhibit complement activating factors (28), blood was also drawn into 15 ml conical tubes without additions and clotted at room temperature for 1 hr. In most cases serum samples were recentrifuged for 5 min to remove residual red blood cells. Plasmas and sera were stored at -20°C until ready for use. Plasma from citrated blood, designated whole plasma (WP), was active when stored at 4°C for over one week. Plasmas were also incubated at 56°C for 30 min to inactivate complement to generate heat-inactivated plasma (HIP) and were likewise stored at -20°C. Normal human serum and sera depleted of C3, C1q, and Factor B were purchased from Complement Technologies.

### Immunoglobulin depletion

WP and HIP were mixed with Protein A-agarose (GoldBio), Protein G-agarose (Invitrogen), and anti-IgM-agarose (Sigma, A9935) at an approximate ratio of 1 volume of plasma to 1 volume of packed beads. The mixtures were rotated for 1 hour, pelleted by centrifugation and depleted supernatants removed from the bead pellet.

### Opsonization reactions

Washed *Mab* (∼1 × 10^7^) were incubated with 33% plasmas, serum, or complement-depleted sera for 20 min at 37°C (26), then diluted in PBS to 12% plasma. The final plasma concentration was 1.2% when added to cells. In some experiments, mannan (1 mg/ml), GlcNAc (100 mM), Mg^2+^/EGTA (5 mM/10 mM), or EDTA (10 mM) was added to bacteria before addition of WP. Opsonized *Mab* were briefly sonicated and added directly to neutrophils. For Western blotting, cells (∼1 × 10^8^) were opsonized for 30 min in 33% plasmas and washed 3 times in PBS before solubilization in SDS sample buffer. In some cases, the supernatant was retained for analysis. Plasmas and sera used were from multiple 5-donor pools.

### Killing assay

Neutrophils were suspended in RPMI supplemented with 10 mM HEPES, pH7.4, and 1-2% pooled heat-inactivated platelet-poor plasma (complete RPMI). Sample tubes contained cells (1×10^6^) suspended in complete RPMI and the respective bacteria at a multiplicity of infection (MOI) of approximately 1:1 in a 0.1 ml volume. Tubes were initially centrifuged at 2000 *g* for 1 min and pelleted cells incubated at 37°C for 5 min to promote bacteria-neutrophil interaction, followed by resuspension and incubation for up to 2 hr, as indicated (27). These experimental conditions promote synchronous, close contact of mycobacteria and neutrophils (57). In some experiments 10% WP and HIP were added to neutrophils immediately before infection with non-opsonized *Mab*. Inhibitor experiments were performed by addition of mannan (1 mg/ml), GlcNAc (100 mM), GalNAc (100 mM), and EDTA (10 mM) immediately before bacterial inoculum. Triton X-100 (0.1%) in 0.9% NaCl was added to inactivate neutrophils and aid in mycobacterial dispersion. Following vortex mixing and serial dilutions in saline, samples were plated on 7H10 agar supplemented with OADC and incubated at 37°C. Colonies were counted 3 to 5 days after plating, and compared to colony counts at initiation of infection. For cell-free killing bacteria were incubated with WP and HIP as for the opsonization reactions, added to complete medium for the indicated times, and enumerated as described above.

### Complement receptor blockade

Neutrophils were pre-incubated for 15 min with 10 μg/ ml anti-CR1 (GeneTex, GTX44217), 30 μg/ml anti-CR3 (BioLegend, clone M1/70, #101248), or the combination. Isotype antibody (BioLegend, #400644, IgG2b) was used in all experiments so that equal protein was present in all experiments.

### Western blotting

Opsonized *M. abscessus* in SDS sample buffer were heated at 95°C for 5 min and loaded on 10% polyacrylamide gels. The gels were transferred to nitrocellulose and *Mab*-associated proteins were detected with anti-C3 (BioLegend, 846302), -C1q (Novus, NB100-64420), -MBL2 (abcam, ab189856), -IgG (abcam, ab97175), -IgA-HRP (Southern Biotech, 2050-05), and -IgM-HRP (Southern Biotech, 2020-05), by enhanced chemiluminescence.

### Statistics

Statistical comparisons were performed by t-test after confirmation of normal distributions.

## Acknowledgements

This research was funded in part by grants from the Cystic Fibrosis Foundation to K.C.M. (MALCOL19I0), J.A.N. (NICK18P0, NICK17K0), T.M.L-P. (LENHAR20D0), and K.S. (and was supported by resources provided by the CFF National Resource Center (NICK20Y2-OUT) and the NIH (R01HL146228).

## Notes

### Competing Interest Statement

The authors have declared no competing interest.

## References

1. Nick JA, Daley CL, Lenhart-Pendergrass PM, Davidson RM. 2021. Nontuberculous mycobacteria in cystic fibrosis. Curr Opin Pulm Med 27:586–592.

2. Victoria L, Gupta A, Gomez JL, Robledo J. 2021. Mycobacterium abscessus complex: A Review of Recent Developments in an Emerging Pathogen. Front Cell Infect Microbiol 11:659997.

3. Sepulcri C, Vena A, Bassetti M. 2023. Skin and soft tissue infections due to rapidly growing mycobacteria. Curr Opin Infect Dis doi:10.1097/QCO.0000000000000905.

4. Martiniano SL, Sontag MK, Daley CL, Nick JA, Sagel SD. 2013. Clinical Significance of a First Positive Nontuberculous Mycobacteria Culture in Cystic Fibrosis. Ann Am Thorac Soc 11:36–44.

5. Nick JA, Dedrick RM, Gray AL, Vladar EK, Smith BE, Freeman KG, Malcolm KC, Epperson LE, Hasan NA, Hendrix J, Callahan K, Walton K, Vestal B, Wheeler E, Rysavy NM, Poch K, Caceres S, Lovell VK, Hisert KB, de Moura VC, Chatterjee D, De P, Weakly N, Martiniano SL, Lynch DA, Daley CL, Strong M, Jia F, Hatfull GF, Davidson RM. 2022. Host and pathogen response to bacteriophage engineered against Mycobacterium abscessus lung infection. Cell 185:1860–1874 e1812.

6. Daley CL, Iaccarino JM, Lange C, Cambau E, Wallace RJ, Andrejak C, Bottger EC, Brozek J, Griffith DE, Guglielmetti L, Huitt GA, Knight SL, Leitman P, Marras TK, Olivier KN, Santin M, Stout JE, Tortoli E, van Ingen J, Wagner D, Winthrop KL. 2020. Treatment of Nontuberculous Mycobacterial Pulmonary Disease: An Official ATS/ERS/ESCMID/IDSA Clinical Practice Guideline. Clin Infect Dis 71:905–913.

7. Bryant JM, Grogono DM, Rodriguez-Rincon D, Everall I, Brown KP, Moreno P, Verma D, Hill E, Drijkoningen J, Gilligan P, Esther CR, Noone PG, Giddings O, Bell SC, Thomson R, Wainwright CE, Coulter C, Pandey S, Wood ME, Stockwell RE, Ramsay KA, Sherrard LJ, Kidd TJ, Jabbour N, Johnson GR, Knibbs LD, Morawska L, Sly PD, Jones A, Bilton D, Laurenson I, Ruddy M, Bourke S, Bowler IC, Chapman SJ, Clayton A, Cullen M, Daniels T, Dempsey O, Denton M, Desai M, Drew RJ, Edenborough F, Evans J, Folb J, Humphrey H, Isalska B, Jensen-Fangel S, Jonsson B, Jones AM, et al. 2016. Emergence and spread of a human-transmissible multidrug-resistant nontuberculous mycobacterium. Science 354:751–757.

8. Honda JR, Virdi R, Chan ED. 2018. Global Environmental Nontuberculous Mycobacteria and Their Contemporaneous Man-Made and Natural Niches. Front Microbiol 9:2029.

9. Howard ST, Rhoades E, Recht J, Pang X, Alsup A, Kolter R, Lyons CR, Byrd TF. 2006. Spontaneous reversion of Mycobacterium abscessus from a smooth to a rough morphotype is associated with reduced expression of glycopeptidolipid and reacquisition of an invasive phenotype. Microbiology 152:1581–1590.

10. Bernut A, Herrmann JL, Kissa K, Dubremetz JF, Gaillard JL, Lutfalla G, Kremer L. 2014. Mycobacterium abscessus cording prevents phagocytosis and promotes abscess formation. Proc Natl Acad Sci U S A 111:E943–952.

11. Byrd TF, Lyons CR. 1999. Preliminary characterization of a Mycobacterium abscessus mutant in human and murine models of infection. Infect Immun 67:4700–4707.

12. Catherinot E, Roux AL, Macheras E, Hubert D, Matmar M, Dannhoffer L, Chinet T, Morand P, Poyart C, Heym B, Rottman M, Gaillard JL, Herrmann JL. 2009. Acute respiratory failure involving an R variant of Mycobacterium abscessus. J Clin Microbiol 47:271–274.

13. Rottman M, Catherinot E, Hochedez P, Emile JF, Casanova JL, Gaillard JL, Soudais C. 2007. Importance of T cells, gamma interferon, and tumor necrosis factor in immune control of the rapid grower Mycobacterium abscessus in C57BL/6 mice. Infect Immun 75:5898–5907.

14. Sanguinetti M, Ardito F, Fiscarelli E, La Sorda M, D’Argenio P, Ricciotti G, Fadda G. 2001. Fatal pulmonary infection due to multidrug-resistant Mycobacterium abscessus in a patient with cystic fibrosis. J Clin Microbiol 39:816–819.

15. Hedin W, Froberg G, Fredman K, Chryssanthou E, Selmeryd I, Gillman A, Orsini L, Runold M, Jonsson B, Schon T, Davies Forsman L. 2023. A Rough Colony Morphology of Mycobacterium abscessus Is Associated With Cavitary Pulmonary Disease and Poor Clinical Outcome. J Infect Dis 227:820–827.

16. Rhoades ER, Archambault AS, Greendyke R, Hsu FF, Streeter C, Byrd TF. 2009. Mycobacterium abscessus Glycopeptidolipids mask underlying cell wall phosphatidyl-myo-inositol mannosides blocking induction of human macrophage TNF-alpha by preventing interaction with TLR2. J Immunol 183:1997–2007.

17. Roux AL, Ray A, Pawlik A, Medjahed H, Etienne G, Rottman M, Catherinot E, Coppee JY, Chaoui K, Monsarrat B, Toubert A, Daffe M, Puzo G, Gaillard JL, Brosch R, Dulphy N, Nigou J, Herrmann JL. 2011. Overexpression of proinflammatory TLR-2-signalling lipoproteins in hypervirulent mycobacterial variants. Cell Microbiol 13:692–704.

18. Roux AL, Viljoen A, Bah A, Simeone R, Bernut A, Laencina L, Deramaudt T, Rottman M, Gaillard JL, Majlessi L, Brosch R, Girard-Misguich F, Vergne I, de Chastellier C, Kremer L, Herrmann JL. 2016. The distinct fate of smooth and rough Mycobacterium abscessus variants inside macrophages. Open Biol 6.

19. Kim SW, Subhadra B, Whang J, Back YW, Bae HS, Kim HJ, Choi CH. 2017. Clinical Mycobacterium abscessus strain inhibits autophagy flux and promotes its growth in murine macrophages. Pathog Dis 75.

20. Malcolm KC, Caceres SM, Pohl K, Poch KR, Bernut A, Kremer L, Bratton DL, Herrmann JL, Nick JA. 2018. Neutrophil killing of Mycobacterium abscessus by intra- and extracellular mechanisms. PLoS One 13:e0196120.

21. Malcolm KC, Nichols EM, Caceres SM, Kret JE, Martiniano SL, Sagel SD, Chan ED, Caverly L, Solomon GM, Reynolds P, Bratton DL, Taylor-Cousar JL, Nichols DP, Saavedra MT, Nick JA. 2013. Mycobacterium abscessus Induces a Limited Pattern of Neutrophil Activation That Promotes Pathogen Survival. PLoS One 8:e57402.

22. Ip WK, Takahashi K, Ezekowitz RA, Stuart LM. 2009. Mannose-binding lectin and innate immunity. Immunol Rev 230:9–21.

23. Schiller NL, Friedman GL, Roberts RB. 1979. The role of natural IgG and complement in the phagocytosis of type 4 Neisseria gonorrhoeae by human polymorphonuclear leukocytes. J Infect Dis 140:698–707.

24. Schwartz JT, Barker JH, Long ME, Kaufman J, McCracken J, Allen LA. 2012. Natural IgM mediates complement-dependent uptake of Francisella tularensis by human neutrophils via complement receptors 1 and 3 in nonimmune serum. J Immunol 189:3064–3077.

25. Lutz HU, Nater M, Stammler P. 1993. Naturally occurring anti-band 3 antibodies have a unique affinity for C3. Immunology 80:191–196.

26. Lenhart-Pendergrass PM, Malcolm KC, Wheeler E, Rysavy NM, Poch K, Caceres S, Calhoun KM, Martiniano SL, Nick JA. 2023. Deficient Complement Opsonization Impairs Mycobacterium avium Killing by Neutrophils in Cystic Fibrosis. Microbiol Spectr 11:e0327922.

27. Pohl K, Grimm XA, Caceres SM, Poch KR, Rysavy N, Saavedra M, Nick JA, Malcolm KC. 2020. Mycobacterium abscessus Clearance by Neutrophils Is Independent of Autophagy. Infect Immun 88.

28. Brady AM, Spencer BL, Falsey AR, Nahm MH. 2014. Blood collection tubes influence serum ficolin-1 and ficolin-2 levels. Clin Vaccine Immunol 21:51–55.

29. Davidson RM, Hasan NA, Epperson LE, Benoit JB, Kammlade SM, Levin AR, Calado de Moura V, Hunkins J, Weakly N, Beagle S, Sagel SD, Martiniano SL, Salfinger M, Daley CL, Nick JA, Strong M. 2021. Population Genomics of Mycobacterium abscessus from U.S. Cystic Fibrosis Care Centers. Ann Am Thorac Soc 18:1960–1969.

30. Bhakdi S, Knufermann H, Fischer H, Wallach DF. 1974. Interaction between erythrocyte membrane proteins and complement components. II. The identification and peptide composition of complement components C3 and C4 desorbed from erythrocyte membranes. Biochim Biophys Acta 373:295–307.

31. Laarman AJ, Ruyken M, Malone CL, van Strijp JA, Horswill AR, Rooijakkers SH. 2011. Staphylococcus aureus metalloprotease aureolysin cleaves complement C3 to mediate immune evasion. J Immunol 186:6445–6453.

32. Ueda T, Rieu P, Brayer J, Arnaout MA. 1994. Identification of the complement iC3b binding site in the beta 2 integrin CR3 (CD11b/CD18). Proc Natl Acad Sci U S A 91:10680–10684.

33. Abrahams KA, Besra GS. 2021. Synthesis and recycling of the mycobacterial cell envelope. Curr Opin Microbiol 60:58–65.

34. Liu Y, Endo Y, Iwaki D, Nakata M, Matsushita M, Wada I, Inoue K, Munakata M, Fujita T. 2005. Human M-ficolin is a secretory protein that activates the lectin complement pathway. J Immunol 175:3150–3156.

35. Sung PS, Chang WC, Hsieh SL. 2020. CLEC5A: A Promiscuous Pattern Recognition Receptor to Microbes and Beyond. Adv Exp Med Biol 1204:57–73.

36. Beuth J, Ko HL, Pulverer G. 1988. The role of staphylococcal lectins in human granulocyte stimulation. Infection 16:46–48.

37. Kolbe K, Veleti SK, Reiling N, Lindhorst TK. 2019. Lectins of Mycobacterium tuberculosis - rarely studied proteins. Beilstein J Org Chem 15:1–15.

38. Hartmann P, Becker R, Franzen C, Schell-Frederick E, Romer J, Jacobs M, Fatkenheuer G, Plum G. 2001. Phagocytosis and killing of Mycobacterium avium complex by human neutrophils. J Leukoc Biol 69:397–404.

39. Fidler KJ, Hilliard TN, Bush A, Johnson M, Geddes DM, Turner MW, Alton EW, Klein NJ, Davies JC. 2009. Mannose-binding lectin is present in the infected airway: a possible pulmonary defence mechanism. Thorax 64:150–155.

40. Hayes D, Jr., Shukla RK, Cheng Y, Gecili E, Merling MR, Szczesniak RD, Ziady AG, Woods JC, Hall-Stoodley L, Liyanage NP, Robinson RT. 2022. Tissue-localized immune responses in people with cystic fibrosis and respiratory nontuberculous mycobacteria infection. JCI Insight 7.

41. Malcolm KC, Wheeler EA, Calhoun K, Lenhart-Pendergrass PM, Rysavy N, Poch KR, Caceres SM, Saavedra MT, Nick JA. 2022. Specificity of Immunoglobulin Response to Nontuberculous Mycobacteria Infection in People with Cystic Fibrosis. Microbiol Spectr 10:e0187422.

42. Khandelwal S, Ravi J, Rauova L, Johnson A, Lee GM, Gilner JB, Gunti S, Notkins AL, Kuchibhatla M, Frank M, Poncz M, Cines DB, Arepally GM. 2018. Polyreactive IgM initiates complement activation by PF4/heparin complexes through the classical pathway. Blood 132:2431–2440.

43. Dubois V, Viljoen A, Laencina L, Le Moigne V, Bernut A, Dubar F, Blaise M, Gaillard JL, Guerardel Y, Kremer L, Herrmann JL, Girard-Misguich F. 2018. MmpL8(MAB) controls Mycobacterium abscessus virulence and production of a previously unknown glycolipid family. Proc Natl Acad Sci U S A 115:E10147–E10156.

44. Palcekova Z, Gilleron M, Angala SK, Belardinelli JM, McNeil M, Bermudez LE, Jackson M. 2020. Polysaccharide Succinylation Enhances the Intracellular Survival of Mycobacterium abscessus. ACS Infect Dis 6:2235–2248.

45. Daher W, Leclercq LD, Viljoen A, Karam J, Dufrene YF, Guerardel Y, Kremer L. 2020. O-Methylation of the Glycopeptidolipid Acyl Chain Defines Surface Hydrophobicity of Mycobacterium abscessus and Macrophage Invasion. ACS Infect Dis 6:2756–2770.

46. Zhou M, Todd RF, 3rd, van de Winkel JG, Petty HR. 1993. Cocapping of the leukoadhesin molecules complement receptor type 3 and lymphocyte function-associated antigen-1 with Fc gamma receptor III on human neutrophils. Possible role of lectin-like interactions. J Immunol 150:3030–3041.

47. Galon J, Gauchat JF, Mazieres N, Spagnoli R, Storkus W, Lotze M, Bonnefoy JY, Fridman WH, Sautes C. 1996. Soluble Fcgamma receptor type III (FcgammaRIII, CD16) triggers cell activation through interaction with complement receptors. J Immunol 157:1184–1192.

48. Neth O, Jack DL, Dodds AW, Holzel H, Klein NJ, Turner MW. 2000. Mannose-binding lectin binds to a range of clinically relevant microorganisms and promotes complement deposition. Infect Immun 68:688–693.

49. Polotsky VY, Belisle JT, Mikusova K, Ezekowitz RA, Joiner KA. 1997. Interaction of human mannose-binding protein with Mycobacterium avium. J Infect Dis 175:1159–1168.

50. Kuhlman M, Joiner K, Ezekowitz RA. 1989. The human mannose-binding protein functions as an opsonin. J Exp Med 169:1733–1745.

51. Dumestre-Perard C, Lamy B, Aldebert D, Lemaire-Vieille C, Grillot R, Brion JP, Gagnon J, Cesbron JY. 2008. Aspergillus conidia activate the complement by the mannan-binding lectin C2 bypass mechanism. J Immunol 181:7100–7105.

52. Haapasalo K, Meri S. 2019. Regulation of the Complement System by Pentraxins. Front Immunol 10:1750.

53. Zacho RM, Jensen L, Terp R, Jensenius JC, Thiel S. 2012. Studies of the pattern recognition molecule H-ficolin: specificity and purification. J Biol Chem 287:8071–8081.

54. Dustin ML. 2016. Complement Receptors in Myeloid Cell Adhesion and Phagocytosis. Microbiol Spectr 4.

55. Nakayama H, Kurihara H, Morita YS, Kinoshita T, Mauri L, Prinetti A, Sonnino S, Yokoyama N, Ogawa H, Takamori K, Iwabuchi K. 2016. Lipoarabinomannan binding to lactosylceramide in lipid rafts is essential for the phagocytosis of mycobacteria by human neutrophils. Sci Signal 9:ra101.

56. Young RL, Malcolm KC, Kret JE, Caceres SM, Poch KR, Nichols DP, Taylor-Cousar JL, Saavedra MT, Randell SH, Vasil ML, Burns JL, Moskowitz SM, Nick JA. 2011. Neutrophil extracellular trap (NET)-mediated killing of Pseudomonas aeruginosa: evidence of acquired resistance within the CF airway, independent of CFTR. PLoS One 6:e23637.

57. Malcolm KC. 2018. Measuring Neutrophil Bactericidal Activity. Methods Mol Biol 1809:139–144.

