## Supplemental Figures S1-S5 for "Divergent host innate immune response to the smooth-to-rough *M. abscessus* adaptation to chronic infection"

Suppl Fig 1

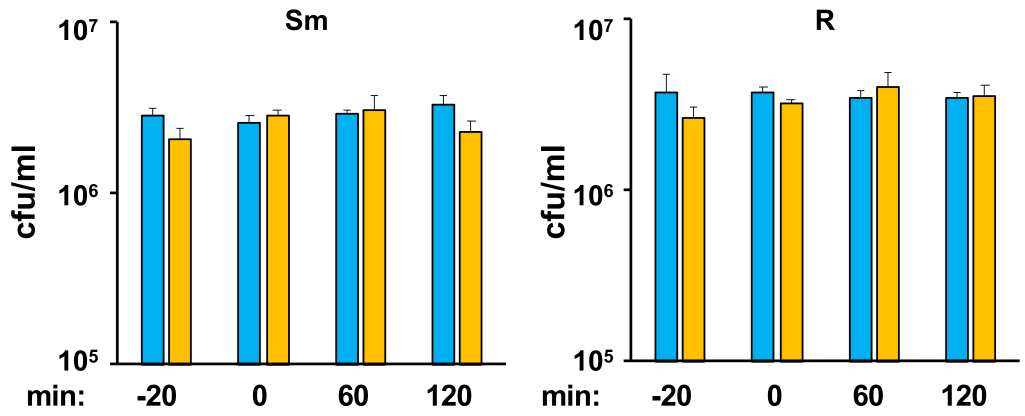


**Supplemental Fig 1. Mab survives opsonization in the absence of neutrophils**. Smooth (Sm) and rough (R) Mab was opsonized with WP (blue) or HIP (orange) alone and incubated for the indicated times. The -20 and 0 times are the beginning and end of the opsonization reaction, respectively ; n= 3. Mab were plated and counted to determine survival.

Suppl Fig 2

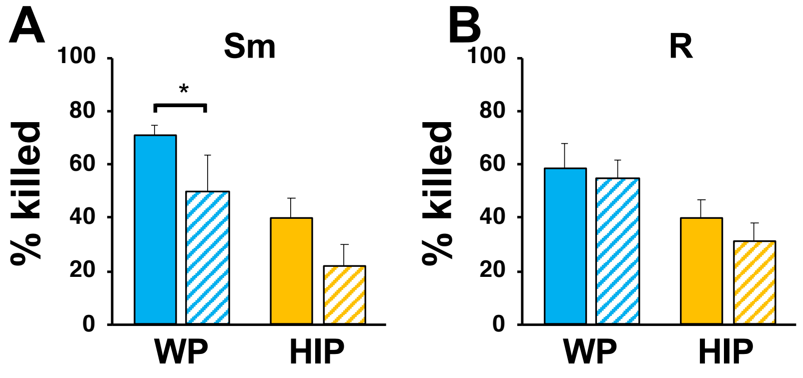


**Supplemental Fig 2. Effect of plasma components in the presence of neutrophils**. **A**, smooth and **B**, rough Mab were pre-opsonized in WP or HIP (closed bars), or non-opsonized Mab was added to neutrophils after addition of 10% WP or HIP (striped bars) and killing determined after 1 hr incubation with neutrophils.

Suppl Fig 3

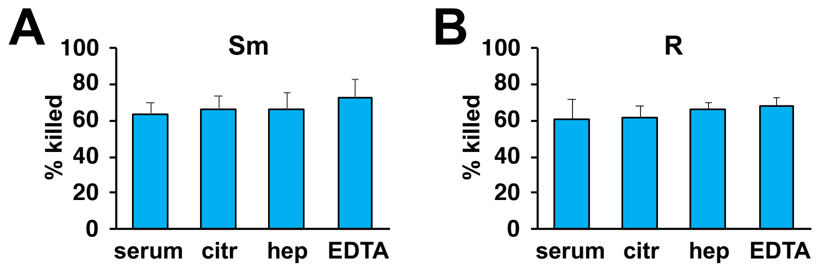


**Supplemental Fig 3. Serum and plasma sources are similarly effective in supporting Mab killing**. **A**, smooth Mab and **B**, rough Mab opsonized with serum, or whole plasmas from blood drawn into citrate, heparin, or EDTA were added to human neutrophils at an MOI of 1 for 1 hr and killing determined; n=4.

Suppl Fig 4

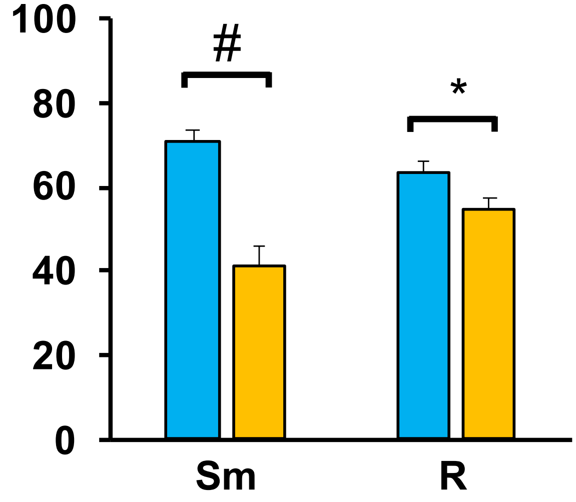


**Supplemental Fig 4. Composite data of killing of smooth and rough Mab clinical isolates**. Data from experiments in Fig 4 were combined to observe the general effect on killing of Mab morphotypes; n = 21 smooth and 42 rough independent experiments. WP (blue) and HIP (orange); *, $P<0.05$; #$P<10^{-5}$.

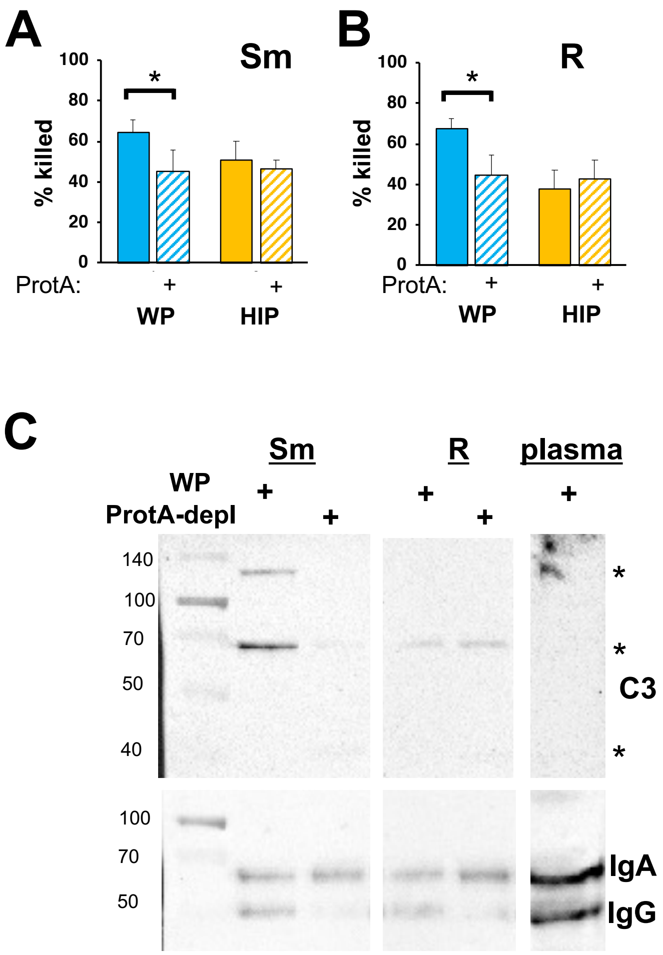


**Supplemental Fig 5. Reduced neutrophil killing of Mab after opsonization with Protein A-depleted WP**. **A**, smooth and **B**, rough Mab opsonized with WP or HIP alone (solid) or in WP or HIP depleted of IgG and IgM with Protein A agarose (striped bars) were added to human neutrophils at an MOI of 1 for 1 hr and killing determined; n= 6. **C**, Mab opsonized as in A were washed, proteins separated by SDS-PAGE, and deposited C3, IgG, and IgA were detected. A 1:20 dilution of WP or Protein A-depleted WP were run as a positive controls. *, P<0.05.
